# Gene essentiality in cancer is better predicted by mRNA abundance than by gene regulatory network-inferred activity

**DOI:** 10.1101/2023.03.02.530664

**Authors:** Cosmin Tudose, Jonathan Bond, Colm J. Ryan

## Abstract

Gene regulatory networks (GRNs) are often deregulated in tumor cells, resulting in altered transcriptional programs that facilitate tumor growth. These altered networks may make tumor cells vulnerable to the inhibition of specific regulatory proteins. Consequently, the reconstruction of GRNs in tumors is often proposed as a means to identify therapeutic targets. While there are examples of individual targets identified using GRNs, the extent to which GRNs can be used to predict sensitivity to targeted intervention in general remains unknown. Here we use the results of genome-wide CRISPR screens to systematically assess the ability of GRNs to predict sensitivity to gene inhibition in cancer cell lines. Using GRNs derived from multiple sources, including GRNs reconstructed from tumor transcriptomes and from curated databases, we infer regulatory gene activity in cancer cell lines from ten cancer types. We then ask, in each cancer type, if the inferred regulatory activity of each gene is predictive of sensitivity to CRISPR perturbation of that gene. We observe slight variation in the correlation between gene regulatory activity and gene sensitivity depending on the source of the GRN and the activity estimation method used. However, we find that there is consistently a stronger relationship between mRNA abundance and gene sensitivity than there is between regulatory gene activity and gene sensitivity. This is true both when gene sensitivity is treated as a binary and a quantitative property. Overall, our results suggest that gene sensitivity is better predicted by measured expression than by GRN-inferred activity.

## Introduction

A large volume of cancer molecular profiles have become available through compendia such as The Cancer Genome Atlas (TCGA) and the Cancer Cell Line Encyclopedia (CCLE)^1^. Additionally, maps of cancer vulnerabilities have been generated using CRISPR and drug screens through efforts such as The Cancer Dependency Map (DepMap) and the Genomics of Drug Sensitivity in Cancer (GDSC)^2–7^. A major outstanding challenge is to identify therapeutic targets for molecularly defined cohorts.

Many genetic alterations drive oncogenesis by altering transcriptional programs that govern critical cellular processes such as proliferation, cell cycle and apoptosis via gene regulatory networks (GRNs)^8^. In cancer, GRN perturbation disrupts key transcriptional programs, and can lead to changes in response or resistance to therapies^9,10^. Targeting GRNs to restore normal cell function is a clinically attractive idea. Currently, there are ongoing trials targeting molecular networks, such as STAT3/5 or menin in acute myeloid leukemia (AML), estrogen receptor in ER+/HER2-breast cancer and MDM2 as part of the p53-MDM2 interaction in various cancer types^11^.

Computational tools are often employed to reconstruct GRNs with a view to identifying therapeutic targets^12,13^, e.g., ARACNe^14^, GENIE3^15^ and KBoost^16^ interpret correlations from transcriptomes to construct GRNs. Each GRN is composed of regulons and each regulon contains a regulatory gene, its targets, and the weights between. These GRNs, however, are *in-silico* inferred, and biological validation is not straightforward. This is particularly challenging in human cells, as a gold standard map of human GRNs does not yet exist.

Transcription factor (TF) activity, also referred to as protein activity^17^ or regulon activity^18^, represents the inferred activity of a regulatory gene derived from the variance in transcript abundance of its targets, according to a pre-determined regulon^19^. Inferred activity has been used to investigate drug response^20,21^, uncover “hidden” drivers^22^ and showcase the role of “master regulators” in cancer^17,23,24^. However, validation often involves assessing the impact of perturbing a small number of example genes and measuring the resulting transcriptional changes^17^. Akin to GRN inference methods, there are many activity inference methods, with little consensus across them^25^.

Given that GRNs have been suggested to drive oncogenic processes, dysregulation of regulatory gene activity may lead to vulnerability to perturbation and dependency to regulatory genes^26,27^. However, this has not been assessed at a systematic level. Here, we used CRISPR screens as a precise method to validate whether GRN-inferred activity can predict sensitivity to inhibition. In CRISPR screens, each gene is perturbed with sgRNAs, and a gene’s sensitivity to inhibition is assigned a score calculated from cell growth and survival^28^. The DepMap project uses this approach to characterize the gene sensitivity profiles of more than 1,000 cell lines via genome-wide CRISPR screens^2–5^. We inferred regulatory gene activity in these cell lines using both computationally derived and curated regulons^29^. We then evaluated correlations between gene sensitivity and inferred activity across cell lines. Additionally, in regulatory genes, we compared expression and activity in their ability to predict sensitivity to inhibition. Overall, we found little evidence of activity estimation methods providing an advantage over measured mRNA abundance.

## Materials and methods

### ARACNe regulons processing

We loaded cancer type-specific ARACNe regulons from the aracne.networks 1.20.0 R package and transformed them into data frames using the ‘reg2tibble’ function from the binilib 0.2.0 R package. We converted Entrez IDs into gene symbols via the org.Hs.eg.db 3.14.0 R package. We calculated an updated mode of regulation (MOR) by multiplying the likelihood with the sign of the MOR for each interaction. MOR indicates the directionality of the interaction (i.e., −1 = inhibition; 1 = activation). ‘Source’ genes missing in the expression data were filtered out.

### GRNdb regulons processing

We downloaded TCGA-inferred cancer type-specific regulons from http://www.grndb.com^30^. We used the gene symbols provided. These regulons contain a weight calculated with GENIE3^15^, but do not contain directionality (inhibition or activation), which the ARACNe regulons do, by providing MOR. Therefore, we inferred MOR using the TCGA dataset, as it was used to build the regulons, as follows:

We downloaded log2(tpm+1) normalized RNA-Seq from the UCSC Treehouse Public Data, v11 Public PolyA. We separated the transcriptomics into ten matrices, for each cancer type, using the “disease” column from the clinical data file. For each cancer type, we removed genes with 0s in more than half of the samples, whilst we imputed the others using “impute.knn” from the impute 1.68.0 R package. We inferred the MOR for each interaction in the GRNdb regulons by calculating the Spearman correlation between the expression of each regulatory gene and its target. We computed an updated MOR by multiplying the GENIE3 weight with the sign of the MOR from the previous step.

### DoRothEA regulons processing

We loaded the human DoRothEA regulons from the dorothea 1.6.0 R package^29^. For the downstream analysis we used the high confidence regulons: A, B and C.

### Data wrangling

We downloaded CCLE gene expression, gene sensitivity data and cell line information from DepMap release 21Q4^2–5^. We filtered these for cell lines from cancer types present in Fig. 1B and for the downstream analysis we kept only cell lines present in both gene expression and gene sensitivity datasets. Likewise, only genes present in both datasets were retained for each cancer type. For gene nomenclature we used HGNC symbols, discarding Entrez IDs. For each cancer individually, we dropped genes with more than 20% 0s across samples in the gene expression profile.

**Fig. 1.**
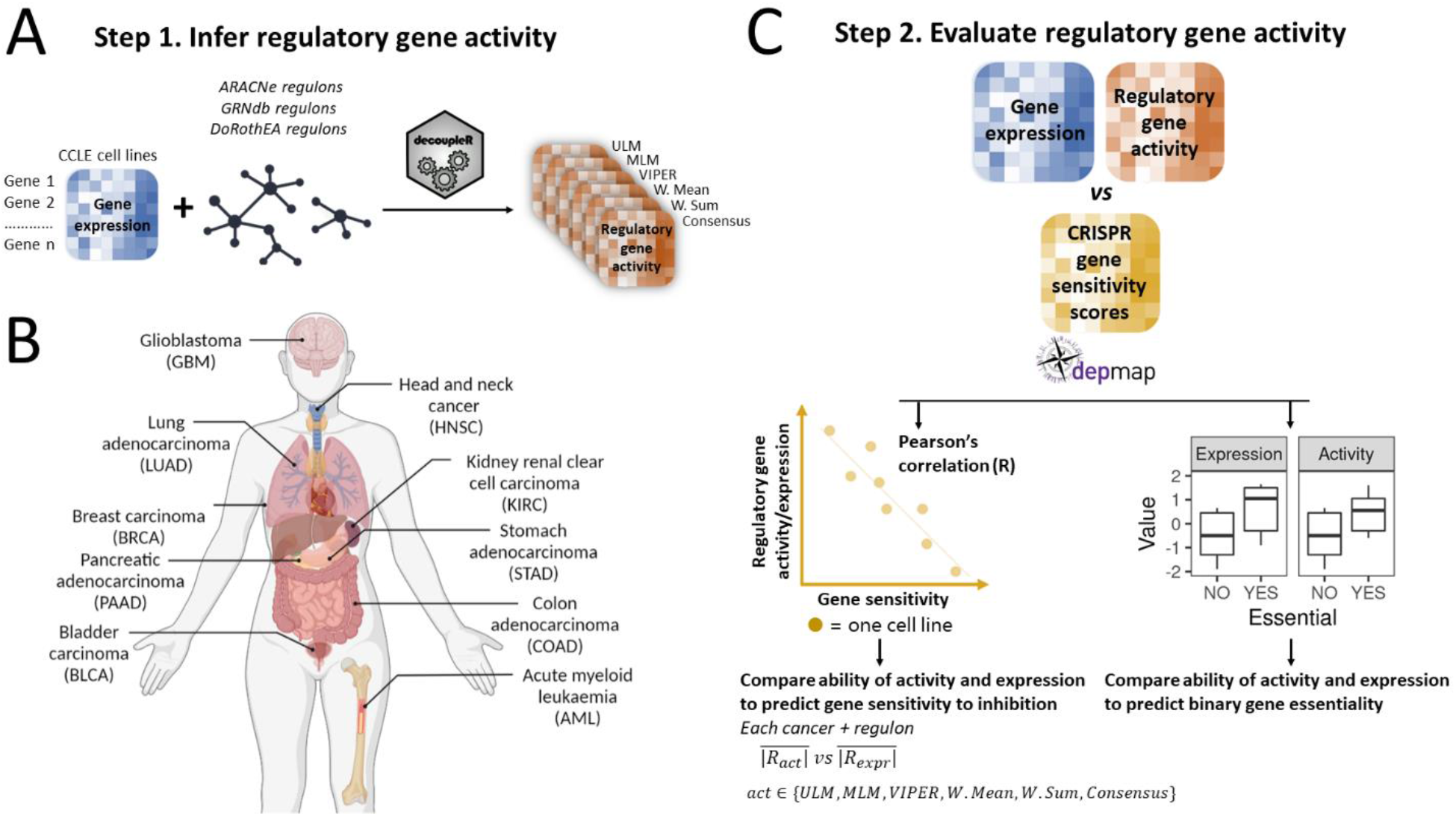
Workflow for evaluating TF activity estimation using CRISPR gene sensitivity profiles from DepMap. **A**, The activity of regulatory genes was inferred in cancer cell lines using six methods from the DecoupleR package. Gene expression profiles from the CCLE were paired with cancer type-specific regulons from ARACNe, GRNdb and curated pan-cancer regulons from DoRothEA to infer regulatory gene activity. **B**, Ten different cancer types (TCGA abbreviations in brackets) with CCLE gene expression profiles and regulons were used for the analysis (generated using BioRender). **C**, Activity inferred using different GRNs and different activity estimation methods was compared with gene expression using two approaches 1. The Pearson’s correlation between inferred activity and gene sensitivity was compared with the Pearson’s correlation between expression and gene sensitivity. 2. Activity and expression were used to look at the degree of separation between essential and non-essential genes in a binary fashion using the common-language effect size (CLES) and a Wilcoxon test.

### Filtering ‘sometimes’ essential genes

We defined genes as essential in a given cell line if they had a CHRONOS score < −0.6 in that cell line. Within each cancer type, we restricted our analysis to genes that were variably essential across cell lines from that cancer type (i.e., genes that were either essential in all cell lines or non-essential in all cell lines were filtered out). Therefore, we were left with sometimes-essential genes only, in order to study variation in sensitivity to inhibition across tumor cells.

For the analysis across all cancers (pan-cancer) we set the threshold to 1% (i.e., nine cell lines). This means for a gene to be considered sometimes essential it had to be essential in at least 1% of cell lines and non-essential in at least 1% of cell lines.

### Computing regulatory gene activity

We ran the ‘decouple’ function from the decoupleR^31^ 2.1.8 package individually on each cancer expression matrix paired with each regulon (See Fig. 1). DecoupleR infers activity via five different methods: Univariate Linear Model (ULM), Multivariate Linear Model (MLM), Virtual Inference of Protein-activity by Enriched Regulon (VIPER), Weighted Mean (W. Mean), Weighted Sum (W. Sum) activity. It then calculates a Consensus across all methods. We used ARACNe, GRNdb and DoRothEA regulons.

### Correlation analysis

For each gene we calculated the Pearson’s correlation between inferred regulatory gene activity/expression and gene sensitivity scores for sometimes essential genes.

We filtered for genes with a significant Pearson’s correlation (p < 0.05) and grouped in categories based on absolute Pearson’s R: 0.2, 0.4, 0.6, 0.8, 1 and based on the sign of R: positive or negative. We used the cor.test function from the R stats 4.1.2 package.

### Enrichment analysis

We ran Gene Ontology (GO) term enrichment analysis using the WebGestalt R package, with FDR = 10%.

We used the genes marked as “oncogene” in the cancer gene census (CGC, https://cancer.sanger.ac.uk/census)^32^ to test for oncogene enrichment in the genes where activity is better correlated with sensitivity, using a Fisher’s exact test. Similarly, we tested for master regulator enrichment using the master regulator list from Paull *et al* (2021) in Table S2^33^.

### Calculate per gene variance for each method

For each gene we calculated the variance across all samples, all regulon sources and cancer type + cancer type-matched regulons.

### Comparing activity methods

For each possible cancer type + cancer type-matched regulon combination we calculated the mean Pearson’s correlation across all genes (no p-value filtering) for each activity method and compared.

We fit a linear model using the lm function from the R stats package (v4.1.0): |R| ~ Cancer type + Regulon source + Activity method + No. cell lines + RNA-Seq variance We estimated the percentage of the variance each term in the linear model explains using adjusted R-squared.

### Comparing regulons

For each possible cancer type + regulon combination, we calculated the mean Pearson’s correlation across all genes (no p-value filtering) for each activity method to investigate whether cancer type-matched regulons are more predictive of sensitivity than mismatched regulons. We plotted the absolute mean Pearson’s R for each combination and assigned ranks 1-10 for each cancer. We conducted Unpaired Two-Samples Wilcoxon tests (wilcox.test function from the R 4.1.2 stats package) to compare the ranks of cancer type-matched regulons to the ranks of cancer type-mismatched regulons.

### Common language effect size calculation

For each cancer type + cancer type-matched regulon combination, we calculated the common language effect size (CLES) across sometimes essential genes for each activity method. Here we considered genes essential in at least three cell lines, and non-essential in at least three cell lines as sometimes essential. We used the CLES function from the bmbstats v0.0.0.9001 R package to predict binary essentiality.

We filtered for significance based on the expression/activity difference in non-essential vs. essential genes (Wilcoxon unpaired test p < 0.05) and counted the number of genes for each method with a CLES > 0.7, > 0.8 and > 0.9, respectively.

R code to run analyses is available at: https://github.com/cancergenetics/GRN_activity_corr_essentiality

## Results

### Variation in correlation between activity and gene inhibition sensitivity is driven more by cancer type than activity estimation method

Estimating regulatory gene activity requires a GRN (containing edges between regulatory genes and targets) and a gene expression matrix (quantifying the expression levels of all genes in a set of samples) (Fig. 1A)^31^. Typically, activity estimation methods assign an activity score to a given regulatory gene such that higher expression of the gene’s targets in a given sample is associated with a higher activity score for the regulatory gene in that sample. Here, rather than focusing on a single GRN or single activity estimation method, we assessed three GRN sources and six activity estimation methods (Fig. 1A).

We selected ARACNe^23,34^, GRNdb^15,30^, and DoRothEA^29^ as representatives of different GRN reconstruction approaches – ARACNe is one of the longest-established methods and infers GRNs solely from transcriptomes; GRNdb is more recently developed and uses GENIE3 GRNs inferred from transcriptomes that are further refined with ChIP-Seq data; while DoRothEA contains curated GRNs that incorporate cis-regulatory information from ChIP-Seq peaks, literature curated resources, and TF binding motifs within promoters. For ARACNe and GRNdb, we obtained cancer type-specific GRNs, i.e., a breast cancer GRN was assembled from gene expression profiles of breast cancer samples, while for DoRothEA, only cancer type-agnostic GRNs were available.

Using decoupleR, we estimated activity in the DepMap cancer cell lines using five different methods: ULM, MLM, VIPER, W. Mean, W. Sum as well as a consensus score calculated by decoupleR using all five scores. These methods work in similar ways: they estimate enrichment scores for each regulatory gene based on its number of targets, targets’ expression, and MOR (i.e., inhibition or activation). We then determined the mean absolute Pearson’s correlation 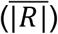 between the regulatory gene activities and CRISPR gene sensitivity scores across cell lines from specific cancer types. Cancer types for which we had matched GRNs derived from relevant TCGA tumor samples were included in this analysis, resulting in ten cancer types being assessed (Fig. 2A). The number of cell lines for each cancer type ranged from 24 (kidney renal clear cell carcinoma - KIRC) to 51 (head and neck squamous cell carcinoma - HNSC) (Fig. 2A). For comparison, we also included 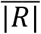 the between mRNA abundance and gene sensitivity scores.

**Figure 2.**
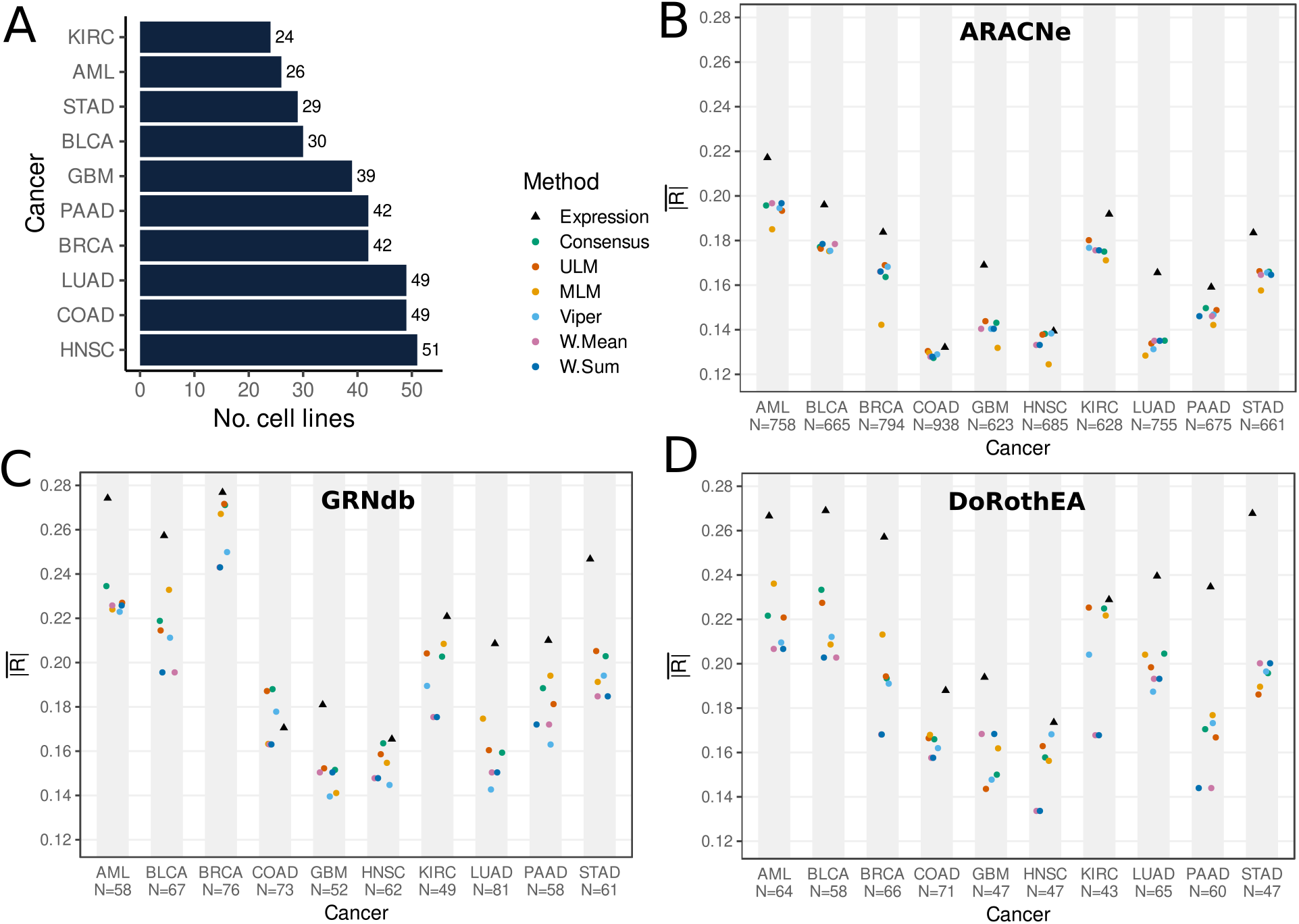
Activity estimation methods have similar performance in predicting gene sensitivity. **A**, Number of cancer cell lines present in DepMap and CCLE for each cancer. **B**, **C, D** Comparison between the different inferred activity methods (paired with cancer type-matched regulons) correlating with gene sensitivity and gene expression correlating with gene sensitivity. **B**, ARACNe. **C**, GRNdb. **D**, DoRothEA (N = number of regulatory genes used to generate 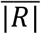 for each cancer type).

We used absolute correlation to assess the association between regulatory gene activity/mRNA abundance and sensitivity because we anticipated that both increased activity (e.g., resulting from amplification) and decreased activity (e.g., resulting from copy number loss) might result in increased sensitivity to inhibition. The former might occur with oncogene addiction-like effects, e.g. *MYC* amplification driving *MYC* sensitivity, while the latter might occur with haploinsufficiency-like effects, e.g. reduced copy number or expression/activity of a gene making cells more sensitive to further perturbation of that gene^35,36^.

Correlations were only calculated for genes that were 1) identified as regulatory genes in the GRN and 2) identified as essential in a subset of cell lines from the cancer type assessed i.e., after excluding genes that are always or never essential (see Methods). A consequence of these criteria is that different GRN methods were evaluated over different gene sets, because they include different regulatory genes (e.g., the ARACNe breast cancer network contains 6,054 regulatory genes, while DoRothEA only contains a total of 271 regulatory genes).

Across all cancer types, the six activity scores yielded an 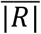 between 0.12 and 0.28 for GRNdb, ARACNe and DoRothEA (Fig. 2B, C, D). Across all cancer types and activity estimation methods, GRNdb shows the highest average correlation (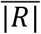 = 0.189), followed by DoRothEA (0.185) and ARACNe (0.156). However, the genes included in each GRN varied significantly – between 628 and 938 genes assessed for ARACNe, between 49 and 81 for GRNdb, and between 43 to 71 for DoRothEA (Fig. 2B, C, D). Summarizing over all regulon sources and cancer types, Consensus had the highest correlation with gene sensitivity, with an 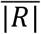 of 0.181. W. Sum and W. Mean performed identically and were jointly the lowest performing of the methods (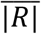 = 0.171).

Although there are differences 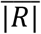 in between the different activity estimation methods and the different GRN sources, visual inspection of the results in Fig. 2 suggests that cancer type may have a much bigger influence than either activity method or GRN source. For instance, although there is variation between the 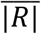 calculated with different activity estimation methods using ARACNe regulons in AML (range 0.18 - 0.2) there is a much bigger difference between the 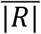 calculated in AML and GBM (median 0.195 for AML; 0.14 for GBM). To understand the relative contributions of different factors to the variability in 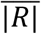, we fit a linear model with three terms: cancer type, regulon source (ARACNe, GRNdb, DoRothEA) and activity method (Consensus, VIPER, etc.). Our results suggest that cancer type does indeed explain 51% of the variance in the model whilst regulon source and activity estimation method explain much less of the variance: 21% and < 0.01%, respectively (Supplementary Fig. S1). Thus, cancer type has a bigger influence than the regulon source, and the activity estimation method shows no consistent contribution.

It is reasonable to ask why cancer type has such a big influence – why would AML cell lines have higher average correlations between activity and gene sensitivity than HNSC cell lines? We reasoned that there might be two explanations: 1) the different numbers of cell lines for each cancer type (ranging from 24 to 51) may result in different distributions of correlations and 2) there may be more transcriptomic diversity in the cell lines from different cancer types. The latter might occur if the cancer in question has more intrinsic heterogeneity or simply if the cell lines available cover more diverse subtypes. We found that there is a strong correlation between 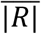 and the number of cell lines used for our analysis (Pearson’s R = −0.89, p < 0.01). In fact, adding the number of cell lines as a variable in the linear model, shows that it explains 29% of the variance when cancer type is excluded, but does not explain any additional variance to cancer type. We find that variance in mRNA abundance together with number of cell lines explained 44% of the variance in the linear model, which is 86% of the variance explained by cancer type (Supplementary Fig. S1). This suggests the majority of the variance explained by cancer type is in fact explained by the number of cell lines analyzed and the variance in mRNA abundance of these cell lines, with unknown factors contributing ~7% in the model.

### Regulons convey cancer type-specific information in relation to gene sensitivity to inhibition

As noted, GRNs for ARACNe and GRNdb are cancer type-specific. We wished to assess whether cancer type-matched GRNs were more informative for predicting gene sensitivity than cancer type-mismatched GRNs. For each cancer type, we ran decoupleR with the cancer type-matched regulons as well as with the nine regulons from the other cancer types (Fig. 1A).

Our results suggest that, on average, for all regulon sources and activity estimation methods, except for MLM, cancer type-matched regulons result in a higher absolute correlation between activity and sensitivity than cancer type-mismatched ones (Unpaired Two-Samples Wilcoxon Test p-value < 0.01) (Fig. 3A, B, Supplementary Table 1). Although cancer type-matched regulons were inferred from patient samples and tested in cell lines, our results suggest that tissue-specific regulon interactions are more relevant, as previously suggested^21^, thereby improving inference of regulatory gene activity and correlation with gene sensitivity in the DepMap. However, despite cancer type-matched regulons performing better than cancer type-mismatched ones, the correlation between activity and gene sensitivity is still relatively poor on average (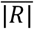 < 0.28).

**Figure 3.**
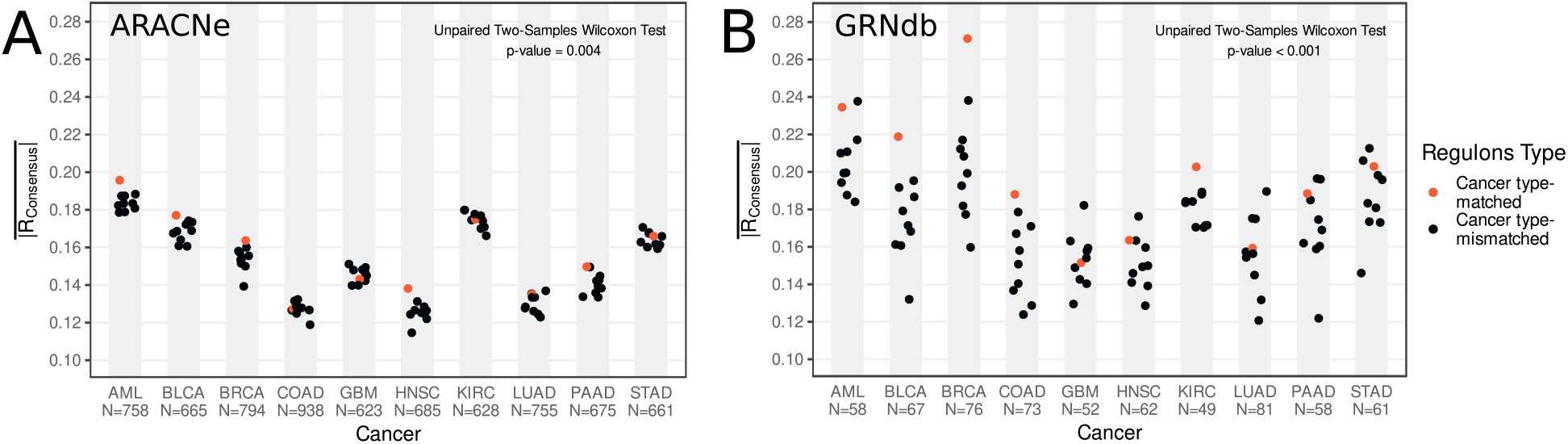
Cancer type-matched regulons predict sensitivity to inhibition better than mismatched regulons. **A, B**, Absolute Pearson correlation between consensus activity and sensitivity for each cancer paired with every regulon. Each dot represents the average absolute Pearson’s correlation coefficients between regulatory gene activity and gene sensitivity across all regulatory genes **A**, ARACNe. **B**, GRNdb. (N = number of regulatory genes used to generate 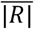 for each cancer type).

### Gene sensitivity to inhibition is better predicted by mRNA abundance than by GRN-inferred activity

We have so far discussed the correlation between regulatory gene activity and gene sensitivity. We have slightly touched on the simpler approach of just using mRNA abundances to predict sensitivity to inhibition. Such a comparison is important for understanding whether the activity estimation methods provide an advantage for predicting gene sensitivity over plain transcript abundance.

Visual inspection of Fig. 2B, C, D suggests that mRNA abundance has a higher correlation with gene sensitivity to inhibition than any of the gene activity estimation methods. This is true across all cancer types analyzed, across all activity estimation methods, and across all GRN sources (Fig. 2B, C, D). While direct comparison of the correlations between regulons from different sources (e.g., ARACNe vs DoRothEA) is challenging due to coverage of different gene sets by each regulon source, this is not the case when comparing the activity estimation methods to mRNA abundance. When comparing the average correlation of activity estimation methods from ARACNe regulons to mRNA abundance we did so over the same set of genes.

We compared the 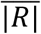 derived from mRNA abundance with that derived from Consensus (the best performing individual activity estimation method) across all regulon sources and all cancer types. We found that mRNA abundance had a significantly higher 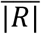 (Wilcoxon paired test p-value = 6.9 × 10^−6^). Overall, this suggests that the average correlation between mRNA abundance and sensitivity to inhibition is higher than that for any of the inferred activity methods using any of the GRNs.

However, for the purpose of identifying new therapeutic targets, strong correlations are more important – those that are highly predictive of gene sensitivity. We therefore compared the proportion of genes that show significant correlations between sensitivity and activity to those with significant correlations between sensitivity and mRNA abundance. We found that strong correlations with gene sensitivity (p < 0.05, |R| > 0.2) were rare. Across all cancer types, neither expression, nor activity had strong correlations with more than 20% of genes (Fig 4B, D, Supplementary Fig. S2A). For nine out of ten cancer types we studied, more genes had a strong correlation between their mRNA abundance and sensitivity than their activity and sensitivity (Fig. 4B, D). KIRC, which has the fewest number of cell lines, was the only cancer type where inferred activity was comparable. Expression consistently had more high correlations than activity across all GRN sources and all activity estimation methods (Fig 4B, D, Supplementary Fig. S2A). The same trend is evident if the threshold for strong correlations is set at (p < 0.05, 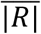 > 0.4) or (p < 0.05, 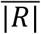 > 0.6). This suggests that, regardless of the exact threshold used to define a strong correlation, gene expression displays more strong correlations with gene sensitivity. Neither GRNdb-inferred activity, nor DoRothEA-inferred activity perform better than mRNA abundance, confirming the findings of ARACNe-inferred activity (Fig 4D, Supplementary Fig. S2A).

**Fig. 4.**
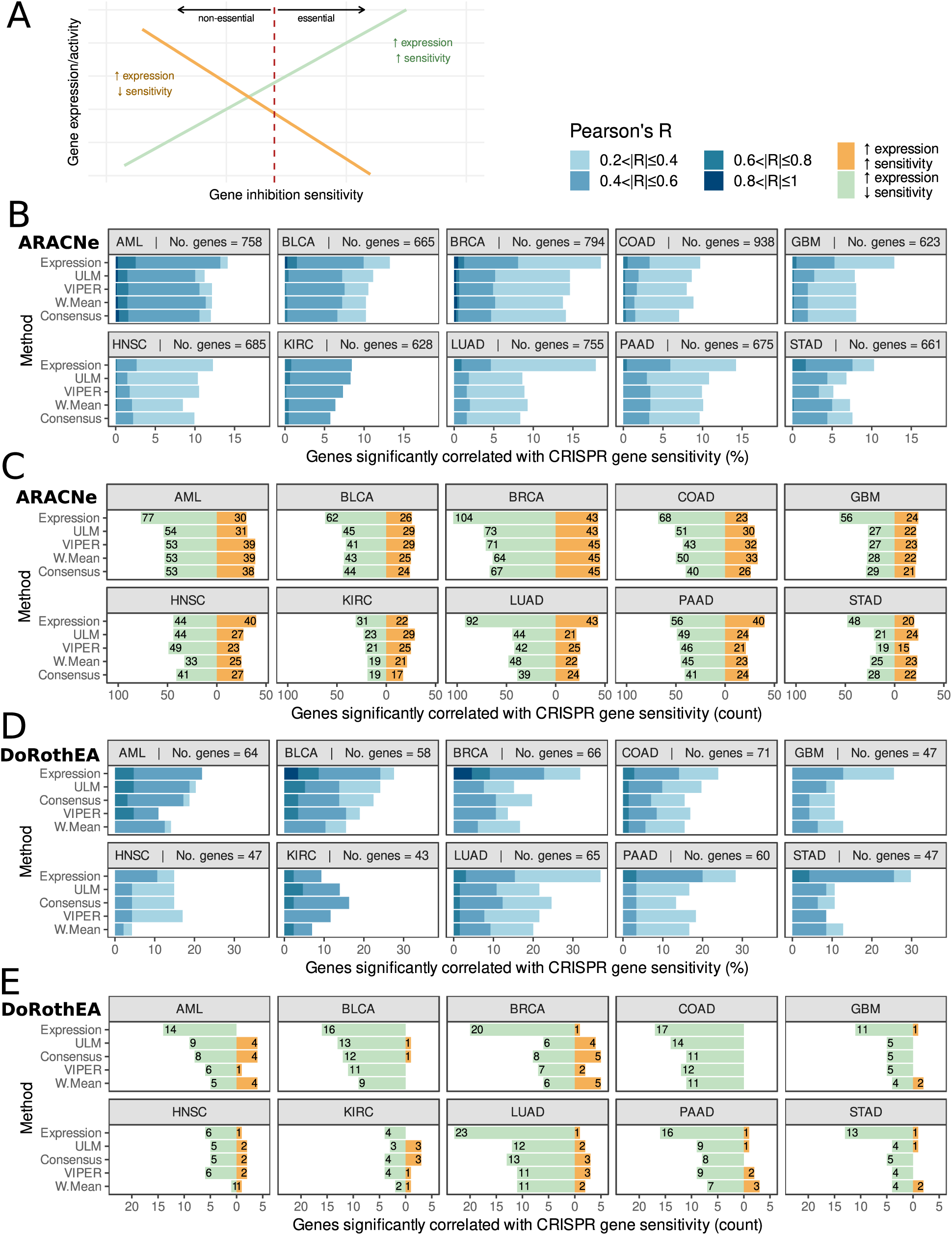
Gene sensitivity to inhibition correlates better with expression than with inferred activity. **A**, Distinction between the two types of correlations between expression/activity and gene sensitivity score. An increase in expression can be correlated with higher sensitivity (in green). An increase in expression/activity can also be associated with a decrease in sensitivity (in orange). **B, D**, Pearson’s correlation coefficients between activity/expression and gene sensitivity stratified incrementally from |R| = 0.2 to 1 to show the percentage of significant regulatory genes correlated with gene sensitivity (p < 0.05) after filtering out genes that are never essential and genes that are always essential in a cancer. Methods are sorted top-to-bottom in order of performance across all cancer types for the GRN-inferred method in cause. **B**, ARACNe regulons. **D**, DoRothEA regulons. **C, E**, Analysis of the positive and negative correlations between activity/expression and gene sensitivity shows there are more negative correlations, suggesting there are more cases where an increase in sensitivity is associated with increased expression/activity. Analysis also confirms expression is better correlated with gene sensitivity. **C**, ARACNe regulons. **E**, DoRothEA regulons.

Whilst expression correlated better with sensitivity to inhibition on average, there were specific cases where inferred activity was a better predictor of sensitivity to inhibition. For example, CDX2 activity performs better in COAD: R_Consensus_ = −0.68, R_Expression_ = −0.49 (Supplementary Table 2). However, we did not find any consistent pattern that explained why these genes were exceptions. For example, GO enrichment analysis did not reveal any specific functional enrichment in genes for which their sensitivity to inhibition is better correlated with activity. Additionally, we found no enrichment for oncogenes among these cases, according to the CGC^32^, or for master regulators, as listed by Paull *et al* (2021)^33^.

To investigate the correlation between activity/expression and sensitivity to inhibition independently of cancer type, we performed the same analysis at a pan-cancer level, using all cell lines (n = 973). We calculated activity based on DoRothEA regulons, as they are cancer type-agnostic. We found a similar trend: ~50% of sometimes-essential genes having a correlation > 0.2 between mRNA abundance and sensitivity to inhibition (Supplementary Fig. S2C). The activity inference methods have strong correlations with fewer genes (25/90 genes with |R| > 0.2 for Consensus vs 43/90 for mRNA abundance). Additionally, expression found two extremely high correlations (|R| > 0.8), whilst activity found none.

One potential explanation for gene expression having higher absolute correlations with gene sensitivity could be that expression measurements display higher variance than activity scores. However, comparing the per gene variance across our methods shows that gene activities, as determined by VIPER and ULM, have a comparable variance to expression, while activities determined by W. Sum have a significantly higher variance. W. Mean, MLM and Consensus have a slightly lower variance than expression (Supplementary Fig. S3).

### Increased sensitivity to gene inhibition is more commonly correlated with increased expression, rather than decreased expression

Thus far, we have focused on the analysis of absolute correlations between activity/expression and sensitivity to inhibition. As noted previously, this is because we anticipated there may be two distinct effect types associated with different genes – sometimes increased expression/activity may be associated with increased sensitivity to inhibition, as observed for oncogene addiction effects, while in other cases reduced expression/activity may be associated with increased sensitivity (Fig. 4A). We sought to understand which type of effect was more common, and whether there were differences between inferred activity and gene expression. We found that for gene expression there were consistently more genes where higher expression was associated with increased gene inhibition sensitivity, as previously suggested by other studies (Fig. 4C, E, Supplementary Fig. S2B)^37,38^. This strong skew towards increased expression – increased sensitivity correlations was less evident for the activity methods, e.g., in AML, using ARACNe regulons, 58% of significant genes (53/91) showed an increase in sensitivity with increased Consensus activity. Across the same gene set 72% of significant genes (77/107) showed an increase in sensitivity with increased gene expression.

There was some variation across the different GRN inference methods, ARACNe in general was associated with a much lower proportion of increased activity – increased sensitivity correlations than GRNdb (Fig. 4C, Supplementary Fig. S2B). However, across both regulon sources, increased expression was consistently associated with an increase in sensitivity.

Interestingly, the use of curated regulons from DoRothEA led to very few cases where an increase in expression/activity results in a decreased sensitivity (Fig. 4E, Supplementary Fig. S2D). This suggests that the TFs included in DoRothEA are skewed towards those for which increased activity/expression is associated with increased inhibition sensitivity.

### Expression better predicts binary essentiality

So far, we have analyzed gene inhibition sensitivity from CRISPR screens as a quantitative trait. However, in many cases the results of CRISPR screens are binarized, such that genes are deemed to be either essential or non-essential for survival^37,39,40^. Genes which are essential in a specific context might then be considered as suitable therapeutic targets.

To assess the ability of gene activity and gene expression to predict binary essentiality, in each cell line we separated genes into two groups: essential and non-essential (see Methods) (Fig. 1B). We then compared the ability of expression and inferred activity to separate the two groups using a Wilcoxon test and the CLES. The interpretation of the CLES is equivalent to the area under the receiver operating characteristic curve (AUC ROC) often used to evaluate binary classifiers. The CLES represents the probability that a gene sampled at random from the essential group will have a higher gene activity/expression than a gene sampled at random from the non-essential group^41^. We consider that a gene’s essentiality can be well predicted by expression/activity if CLES > 0.7 and p-value < 0.05.

Our results suggest that, on average, gene expression better predicts binary essentiality, irrespective of whether ARACNe, GRNdb or DoRothEA regulons were used (Fig. 5A, B, C, Supplementary Table 3). In 20 of 30 cases across all regulon types (ten cancer types x three regulon sources), more genes have a CLES > 0.7 when their essentiality is predicted using expression, rather than activity. The same is true for different thresholds – for CLES > 0.8 and > 0.9, expression still predicts more essential genes overall than any of the activity estimation methods using any of the regulon sources. Similarly, on the DoRothEA pan-cancer analysis we found that expression finds ~51% of sometimes-essential genes with a CLES > 0.7 and ~32% of genes with a CLES > 0.9. Consensus, the best performing activity method finds ~28% genes with a CLES > 0.7 (Supplementary Fig. 4). Thus, gene expression, rather than inferred activity, is a better predictor of binary gene essentiality.

**Fig. 5.**
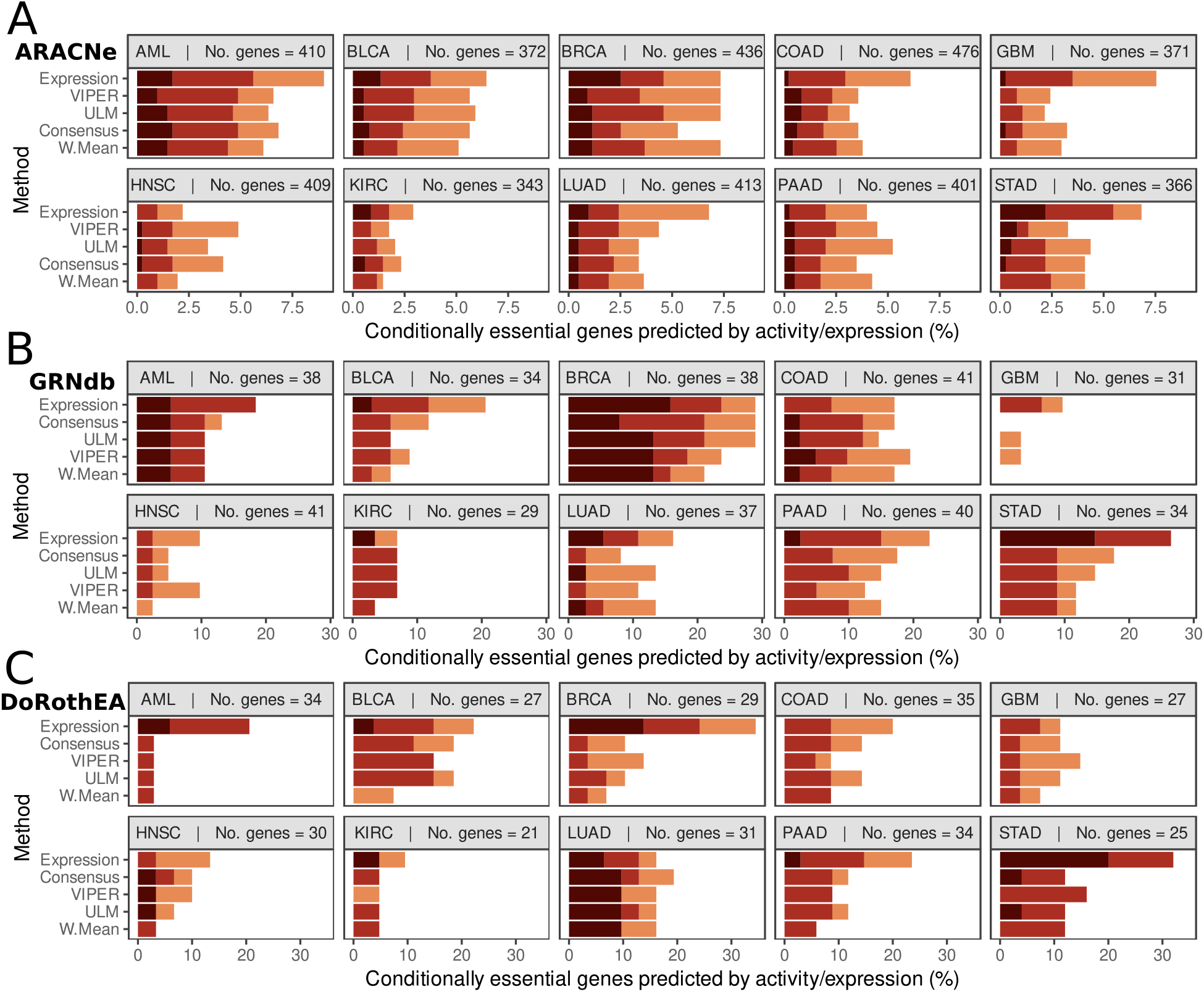
Gene essentiality correlates better with expression than with inferred activity. **A, B, C**, CLES between activity/expression and binary gene essentiality stratified incrementally from CLES = 0.7 to 1 to show the percentage of significant conditionally essential genes predicted by activity/expression (p < 0.05) after filtering out genes that are not essential and genes that are essential in less than three cell lines in a cancer type. Methods are sorted top-to-bottom in order of performance across all cancer types for the GRN-inferred method in cause. **A**, ARACNe regulons. **B**, GRNdb regulons. **C**, DoRothEA regulons.

## Discussion

Our systematic analysis suggests that gene expression performs better than GRN-inferred activity at predicting sensitivity to CRISPR gene inhibition in cancer. This is true regardless of the GRN source and activity inference method used and whether essentiality is treated as a binary or quantitative trait. Whilst extensively used to find “master regulators” of cancer^29^, regulatory gene activity does not outperform gene expression for the task of predicting gene sensitivity to inhibition. Across ten cancer types and at a pan-cancer level, more genes are found to have a strong correlation between sensitivity and mRNA abundance than they do between sensitivity and inferred activity.

We find that matched regulons may be more accurate in describing the regulatory gene activity landscape of each cancer type than mismatched regulons. This suggests that there is value to using co-expression information from relevant patient tumors to build GRNs. Additionally, this suggests a degree of similarity between primary patient tumors and cell-line models that can be captured by GRNs. Our study also suggests that cancer type contributes more to the variance in average correlation with gene sensitivity than the GRN-building method or the activity-inference method. This may be at least partially attributable to different cancer types having more variable transcriptomes. We find that there is no significant difference between activity and expression in finding correlations where an increase in sensitivity is associated with decreased expression/activity. However, in all cancer types, there are more genes where an increase in expression, rather than activity, is associated with an increase in sensitivity, in an oncogene addiction-like effect. This is expected to be the case in most cancer cells reliant on the activity of a TF for survival.

Both inferred and curated networks^29^ have been used to infer the activity of TFs and to investigate the differential activity of TFs in different conditions^21,42^. The assumption behind these approaches is that the activity of a TF inferred from the expression profile of its targets can help uncover hidden potential therapeutic targets. Therefore, these methods have been used as hypothesis-creation tools to find novel targets^21^ or associations with patient survival^43^. A significant limitation of TF activity estimation approaches is that typically only a small number of candidate targets are selected for experimental testing^17^. It is thus extremely challenging to understand how broadly useful these approaches are, and to estimate false positive or false negative rates.

We propose a computational approach based on CRISPR screen data to assess the ability of GRN-inferred activity to predict sensitivity to perturbation in tumor cell lines. The repository of cell lines being screened grows every year, offering more statistical power^2–5^. A significant advantage of our approach is that it is unbiased, in the sense that all genes are evaluated, rather than one or two selected candidates. Evaluating only one or two candidates may lead to a false sense of accuracy of the approach, downplaying its limitations. A limitation of our approach is that we are evaluating a downstream use of inferred GRNs rather than the GRNs themselves, i.e., we have not evaluated the ability of the reconstructed networks to predict transcriptional changes, but rather their ability to predict therapeutic targets. However, the latter is a purpose for which they are often employed.

Our study is primarily limited by the availability of the data, as there is a limited number of cell lines with genome-wide CRISPR screens data. We tried to mitigate this by selecting cancer types with a large number of cell lines screened. Furthermore, the gene sensitivity measurements we use are made in cell line models. These might not entirely reflect cancer cells within actual tumors, surrounded by the tumor microenvironment. Additionally, our results are reflective of cohorts of cell lines displaying a range of regulatory landscapes. A better approach might be to integrate multiple sources of data for model-specific GRNs, as done by Goode *et al*. (2016)^44^ and Assi *et al*. (2019)^45^. They create GRNs from multiple omics sources that are cell line-specific.

Ultimately, our study primarily looks at correlations between GRN-inferred activity and sensitivity and future work could explore indirect relationships between the GRNs and sensitivity to inhibition. For instance, Garcia-Alonso *et al*. (2018)^21^ explores the relationships between drug sensitivity and the activity of indirect targets, finding correlations with clinical significance. However, despite these caveats, expression consistently outperforms GRN-inferred activity in predicting sensitivity to CRISPR inhibition in a variety of cancer types. This work underlines the utility of sensitivity data from CRISPR screens in benchmarking the use of GRN-inferred activity methods for nominating therapeutic targets.

## Acknowledgements

This research was funded by Science Foundation Ireland through the SFI Centre for Research Training in Genomics Data Science under Grant number 18/CRT/6214 and supported in part by the EU’s Horizon 2020 research and innovation programme under the Marie Skłodowska-Curie grant H2020-MSCA-COFUND-2019-945385.

Work in the Bond laboratory is supported by Science Foundation Ireland grants 20/FFP-P/8844 and 18/SPP/3522, the latter together with Children’s Health Ireland. Work in the Ryan laboratory is supported by Science Foundation Ireland grant 20/FFP-P/8641.

We thank Dr. Christina Kiel, Dr. Luis Iglesias Martinez and members of the Bond and Ryan labs for useful suggestions on the analysis. We thank Philip Cotter for technical support on maintaining the server on which many of the scripts were run.

The results shown here are in part based upon data generated by the TCGA Research Network: https://www.cancer.gov/tcga.

---

**Supplementary Table 1.** P-value table of the unpaired two-samples Wilcoxon test comparing the ranks of the activity vs essentiality absolute correlation between cancer type-matched and cancer type-mismatched regulons for each activity method. (Same as in Fig 3A, B).

| Unpaired Two-Samples Wilcoxon Test<br>p-values |  |  |
| --- | --- | --- |
| Method | ARACNe | GRNdb |
| Consensus | 0.004 | <0.001 |
| ULM | 0.006 | <0.001 |
| MLM | 0.206 | 0.002 |
| W. Mean | 0.008 | <0.001 |
| W.Sum | 0.008 | <0.001 |
| VIPER | 0.011 | 0.003 |

**Supplementary Fig. 1.**
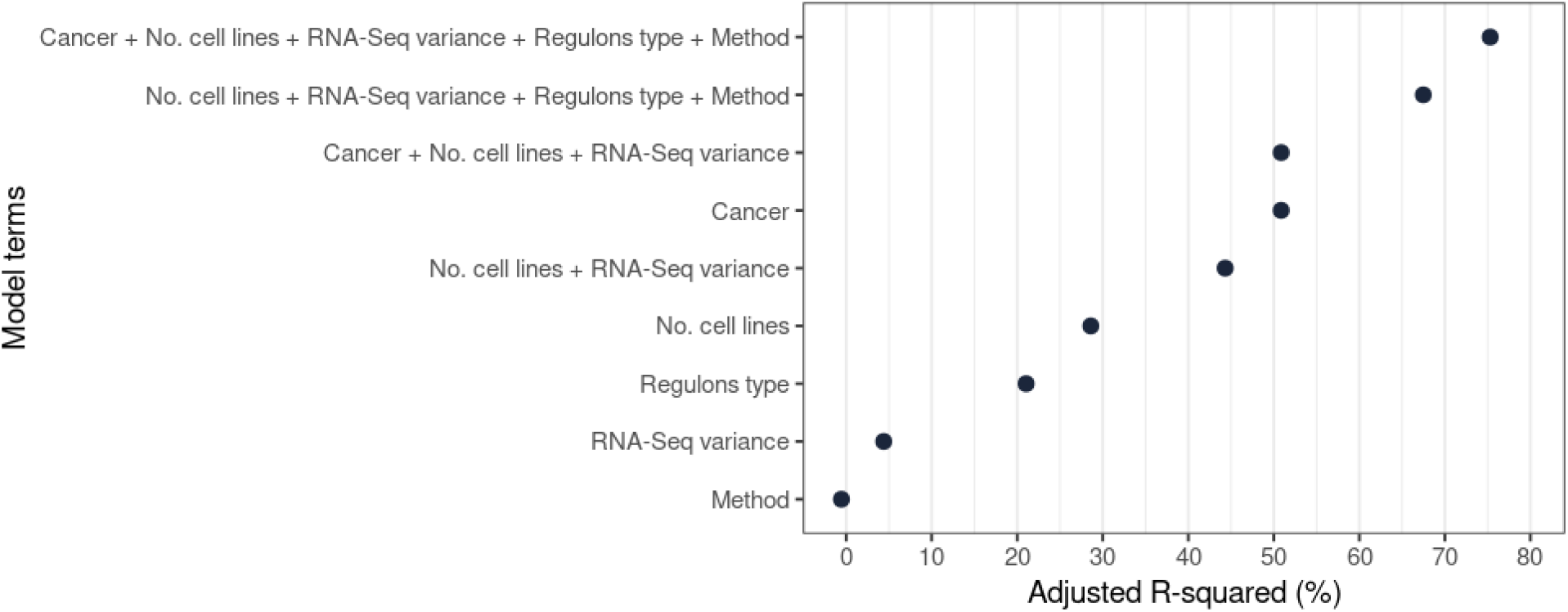
Each terms’ contribution to linear model predicting |R|. Each dot represents the percentage of variance explained (Adjusted R-squared) by each variable in the linear model predicting the absolute correlation between essentiality and activity (|Pearson’s R|).

**Supplementary Fig. 2.**
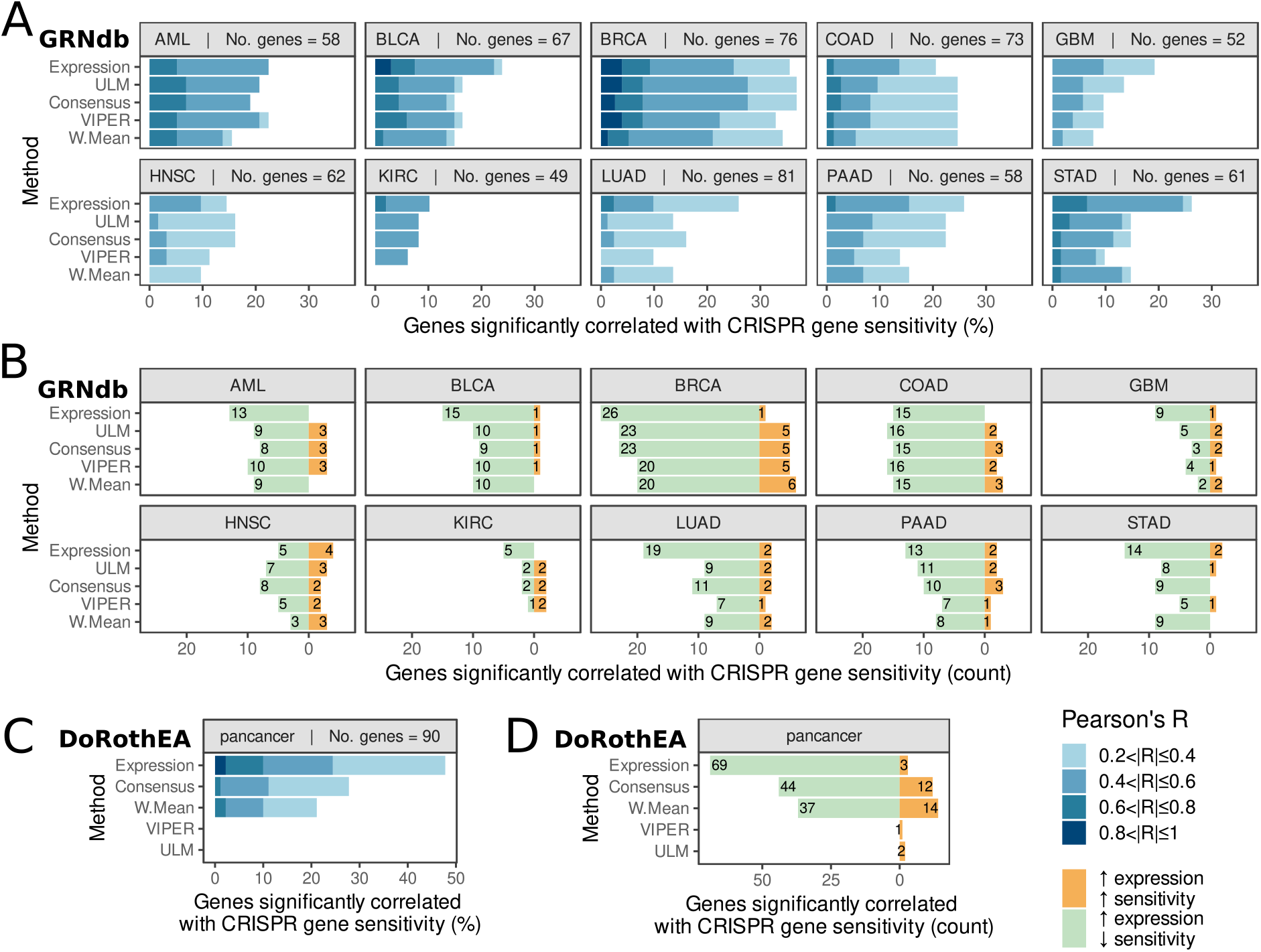
Gene essentiality correlates better with expression than with inferred activity. **A, C**, Pearson’s correlation coefficients between activity/expression and gene sensitivity stratified incrementally from |R| = 0.2 to 1 to show the percentage of significant regulatory genes correlated with gene sensitivity (p < 0.05) after filtering out genes that are never essential and genes that are always essential in a cancer. Methods are sorted top-to-bottom in order of performance across all cancer types for the GRN-inferred method in cause. **A**, GRNdb regulons. **C**, DoRothEA regulons on pan-cancer analysis. **B, D**, Analysis of the high expression – high sensitivity and high expression low sensitivity correlations between activity/expression and gene essentiality shows there are more cases where an increase in sensitivity is associated with increased expression/activity. **B**, GRNdb regulons. **D**, DoRothEA regulons on pan-cancer analysis.

**Supplementary Fig. 3.**
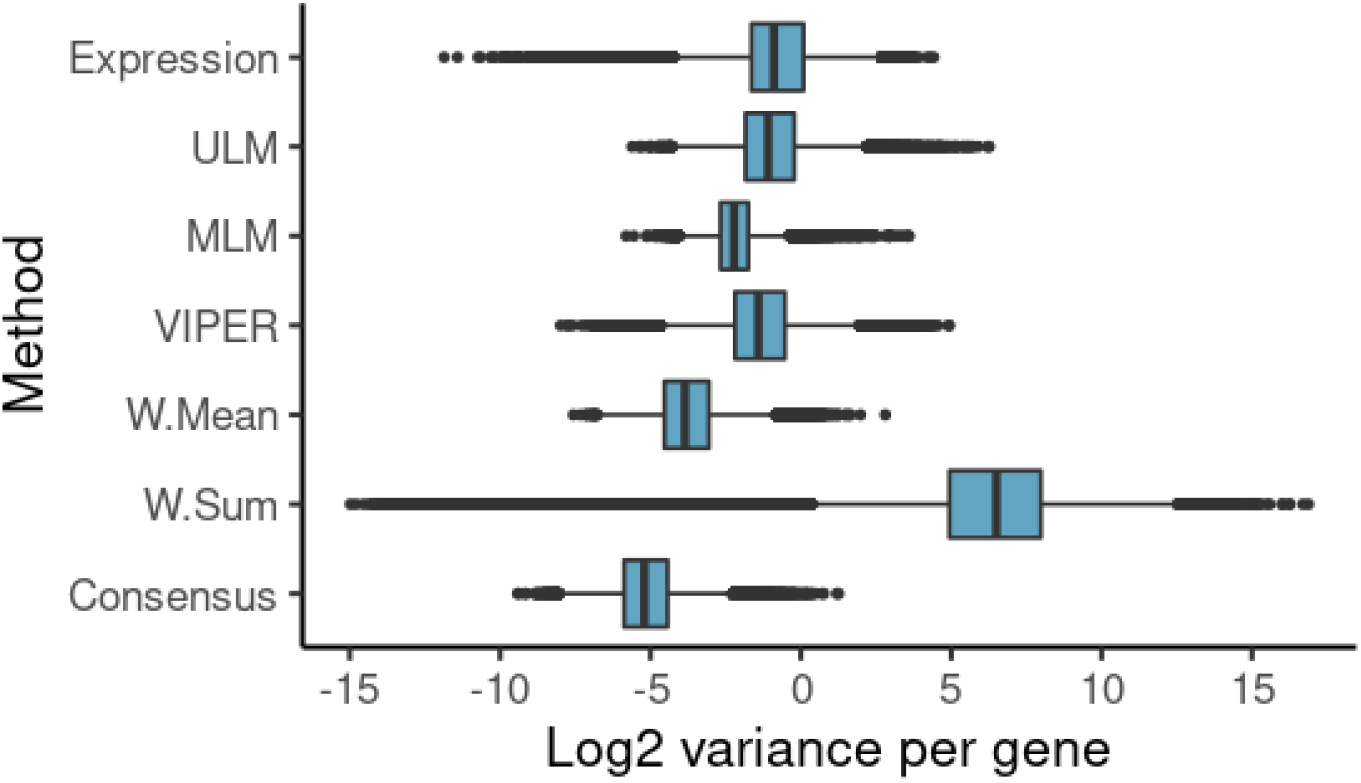
Per gene variance for each method, across all regulon sources. Boxplots showing the log2 variance for each activity method and expression. Black line shows median; blue box represents the interquartile range; black dots show outliers.

**Supplementary Fig. 4.**
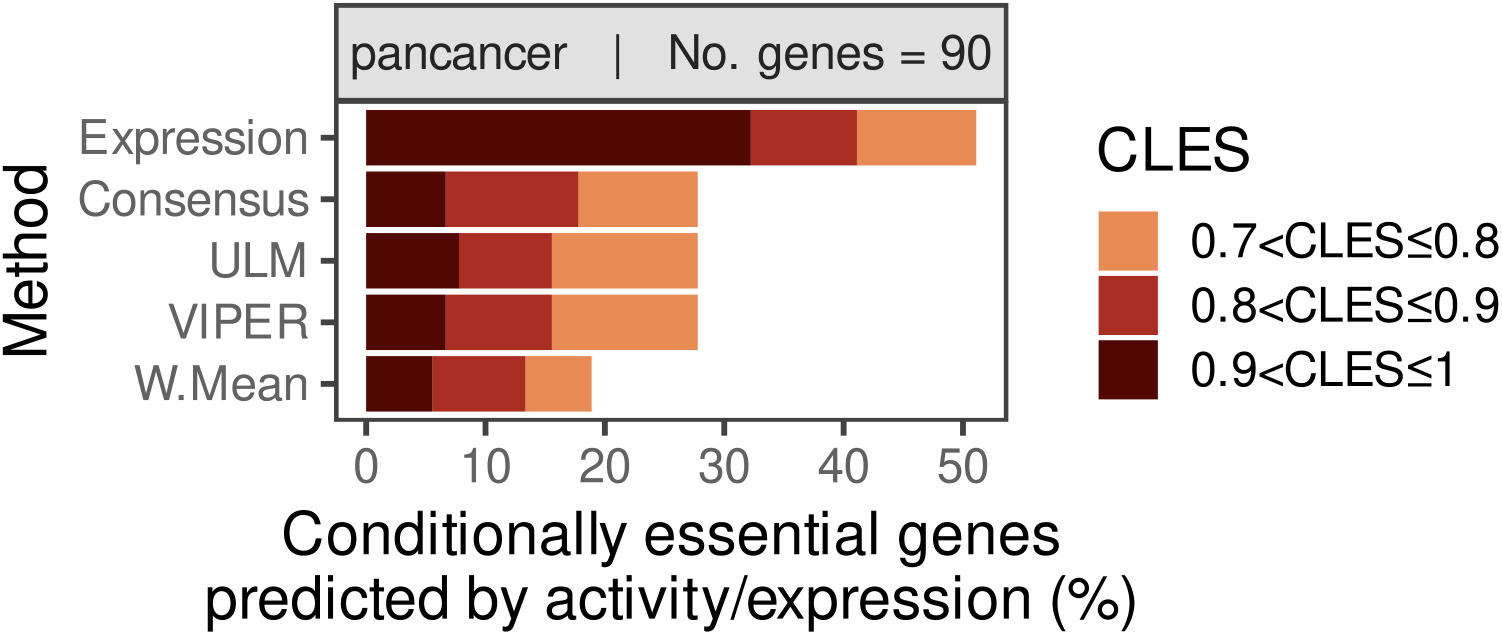
Gene essentiality correlates better with expression than with inferred activity using DoRothEA regulons across in a pan-cancer analysis. CLES between activity/expression and binary gene essentiality stratified incrementally from CLES = 0.7 to 1 to show the percentage of significant conditionally essential genes predicted by activity/expression (p < 0.05) after filtering out genes that are not essential and genes that are essential in less than three cell lines in a cancer type. Methods are sorted top-to-bottom in order of performance across all cancer types for the GRN-inferred method in cause.

## Notes

### Competing Interest Statement

The authors have declared no competing interest.

### Summary of Updates

Figure 4B and 4D were duplicated. Figure 4B should contain results generated with ARACNe, while Figure 4D should contain results generated with DoRothEA.

